# Structural and kinetic insights into a metagenomics-derived Cas12a with high specificity

**DOI:** 10.64898/2026.05.13.724879

**Authors:** Jacquelyn T. Wright, Satya Yendluri, Nicole C. Thomas, Cristina N. Butterfield, Tyler L. Dangerfield, David W Taylor

## Abstract

CRISPR–Cas12a nucleases provide an attractive alternative to Cas9 due to their compact RNA scaffold, T-rich PAM requirement, and improved target specificity. However, the mechanistic features that govern activity and discrimination across Cas12a orthologs remain incompletely understood. Here, we characterize Cas12a-MG29-1, a highly active and specific nuclease identified through metagenomic mining, using cryogenic electron microscopy, mutational analysis, and kinetic modeling. The Cas12a-MG29-1 structure reveals repositioned flexible loops near the distal end of the R-loop, including reduced engagement of one loop region and additional contacts formed by a second distal loop. Structure-guided mutagenesis and loop-swap experiments indicate that distal R-loop architecture modulates target discrimination in a context-dependent manner. Single-turnover cleavage and stopped-flow measurements show that Cas12a-MG29-1 and AsCas12a form reversible R-loops with similar kinetics but differ in strand cleavage following R-loop formation. Global kinetic modeling demonstrates that Cas12a-MG29-1 exhibits accelerated non-target strand cleavage, shifting kinetic partitioning toward product formation. This faster irreversible commitment provides a mechanistic explanation for enhanced activity and specificity without altering initial target interrogation. Together, these findings identify distal R-loop interactions and catalytic commitment as key determinants of Cas12a function and provide a framework for interpreting and engineering next-generation Cas12a orthologs.

## INTRODUCTION

CRISPR (clustered regularly interspaced short palindromic repeats)-Cas (CRISPR associated) protein systems evolved in prokaryotes as a form of adaptive immunity against mobile genetic elements^1–3^ and have been repurposed for use in genome editing and a wide range of other biotechnological applications.^4–6^ Since their initial discovery, a broad diversity of CRISPR systems with distinct activities has been identified and characterized.^7,8^ Among these, Cas9 and Cas12a are the most widely used RNA-guided nucleases for genome engineering applications.

While Cas9 has become the dominant gene-editing platform in both basic and applied research, its broader therapeutic deployment is limited by off-target activity, delivery challenges associated with its relatively large size, and a restricted targeting range imposed by its protospacer adjacent motif (PAM) requirement.^9–12^ Cas12a (previously Cpf1), addresses several of these limitations, as it exhibits higher target specificity,^13^ a slightly more compact effector size and significantly shorter RNA scaffold, and a T-rich PAM,^14^ making it an attractive complement to Cas9. In addition, Cas12a possesses the ability to process its own CRISPR RNA (crRNA),^15–17^ enabling facile multiplexed targeting for applications such as disease modeling^18,19^ and genetic screening.^20,21^ Beyond genome editing, Cas12a also exhibits nonspecific collateral (*trans*) nuclease activity upon target recognition,^22,23^ a property that has been widely exploited for nucleic acid diagnostics.^24–26^ In an effort to expand the known diversity of Cas enzymes, Cas12a-MG29-1 was previously identified through metagenomic mining and shown to exhibit high activity and specificity across multiple cell types.^27,28^ Phylogenetic analysis places Cas12a-MG29-1 within a clade containing well-characterized Cas12a orthologs, including *Francisella novicida* (Fn), *Acidaminococcus sp*. (As), and *Lachnospiraceae bacterium* (Lb) Cas12a, despite substantial sequence divergence from AsCas12a (∼31% amino acid identity; Fig. 1A). However, the mechanistic basis for its high activity and specificity remain unknown.

**Figure 1:**
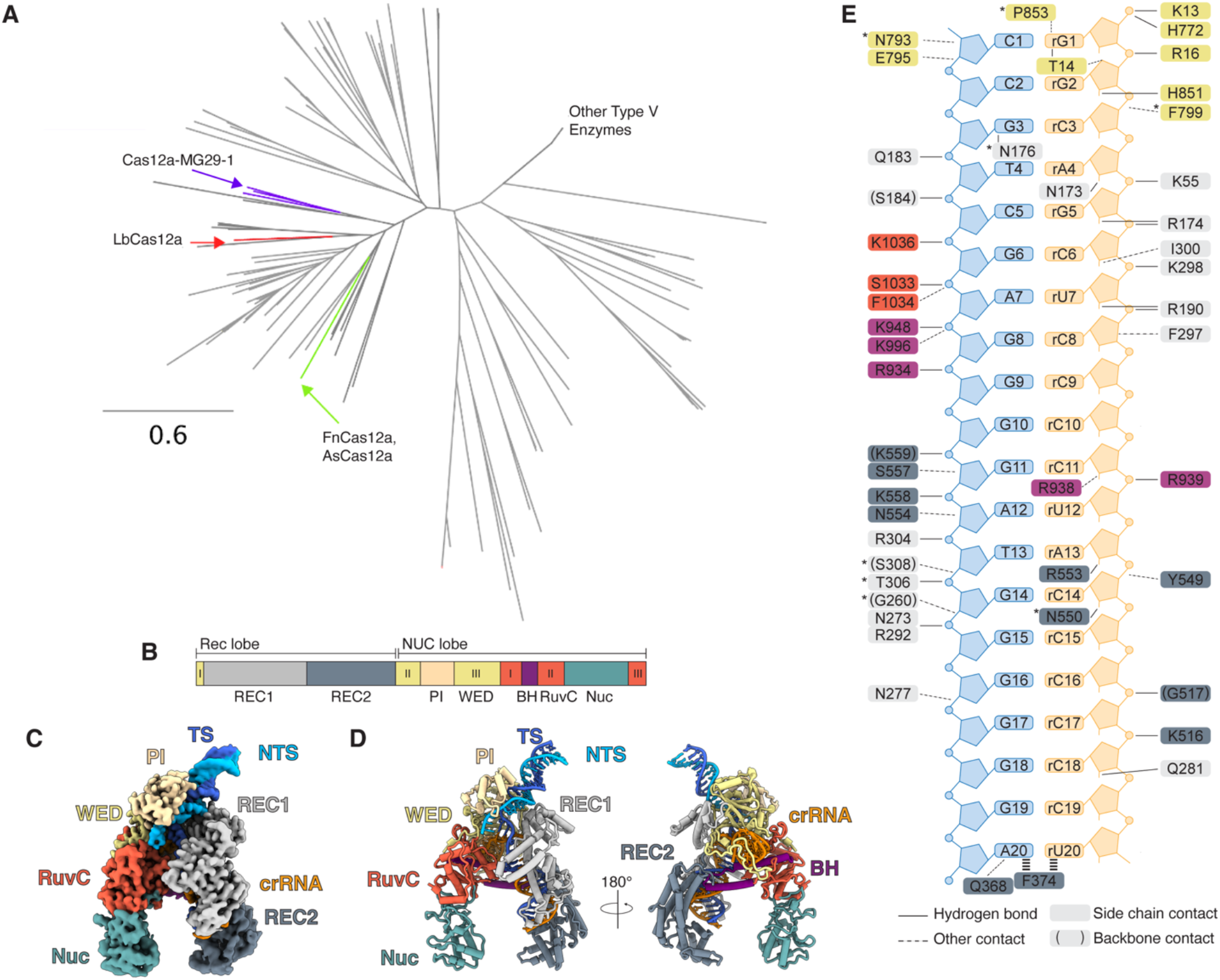
Cas12a-MG29-1 adopts a canonical Cas12a architecture within a conserved phylogenetic lineage. (A) Phylogenetic tree of Cas12a orthologs. Reprinted with permission.^27^ (B) Domain organization of Cas12a-MG29-1. The recognition (Rec) and nuclease (NUC) lobes are shown, including REC1, REC2, PI, WED, bridge helix (BH), RuvC, and Nuc domains. Roman numerals denote discontinuous segments of the WED (I–III) and RuvC (I–III) domains that are separated in sequence but form contiguous structural domains. (C) Front view of post-processed cryo-EM map of the ternary complex of Cas12a-MG29-1. Nucleic acid is colored as follows: crRNA, dark orange; non-target strand, deep sky blue; target strand, royal blue. Protein domains are colored as in (B). (D) Front and back views of the cryo-EM model of the Cas12a-MG29-1 ternary complex, colored according to domain organization in (B) and nucleic acid in (C). (E) Schematic diagram of nucleic acid-protein contacts within the R-loop of the post-cleavage Cas12a-MG29-1 complex. Nucleic acid is colored as in (C) and (D). Protein contacts are colored as in (B). Solid lines indicate hydrogen bonds, while dotted lines denote other contacts. Residues in parentheses refer to side backbone contacts.

Cas12a specificity is enforced during R-loop formation, where a late, rate-limiting transition state acts as a checkpoint for crRNA-target DNA complementarity prior to nuclease activation.^29,30^ During R-loop propagation, few contacts are formed between Cas12a and the RNA-DNA heteroduplex, resulting in a reversible R-loop that enables mismatched targets to destabilize the complex prior to cleavage. As the final base pairs of the R-loop are formed, domain rearrangements facilitate distal protein-nucleic acid contacts, promote nuclease activation, and extend sequence discrimination to late stages of target recognition.^15,30–34^ These observations indicate that protein-nucleic acid interactions near the distal end of the R-loop may play an important role in coordinating catalytic activation with target discrimination. While multiple Cas12a orthologs have been structurally and biochemically characterized, the mechanistic basis for activity and specificity in Cas12a-MG29-1 remains unclear.

Here, we elucidate the mechanistic basis for the high activity and specificity of Cas12a-MG29-1 by combining cryogenic electron microscopy (cryo-EM) with quantitative kinetic analysis. We determined the structure of Cas12a-MG29-1 bound to its crRNA and target DNA and identified differences in protein-nucleic acid contacts near the distal end of the R-loop relative to other Cas12a orthologs. Structure-guided mutagenesis and loop-swap experiments indicate that these regions influence target discrimination without substantially altering global architecture. Single-turnover cleavage and stopped-flow fluorescence measurements show that Cas12a-MG29-1 and AsCas12a form reversible R-loops with comparable kinetics but differ in the rate of cleavage following R-loop formation. Together, these findings provide a framework for understanding how structural and kinetic differences contribute to functional variation among Cas12a orthologs.

## MATERIALS AND METHODS

### Protein Expression and Purification

#### Purification of Cas12a-MG29-1

His-MBP-Cas12a-MG29-1 plasmid was transformed into BL21 (DE3) cells (New England Biolabs). Six colonies were isolated and screened for target protein expression via SDS-PAGE. The colony with the highest expression levels determined by SDS-PAGE was expanded into a working cell bank. A 1L starter culture was grown overnight in Terrific Broth (TB) media containing 50 mg/L kanamycin and 5 g/L glucose anhydrous at 37°C and 185 rpm. 200mL of the overnight culture was added to 1.3L of batch media (55.9 g/L TB media, 0.75 g/L citric acid, 1.4 g/L ammonium sulfate, 1.2 g/L magnesium sulfate, 5 g/L glucose, 50 mg/L kanamycin, 0.2 mL/L antifoam, and 1x trace metals) to a Minifors-2 Bioreactor (Infors-HT) and grown at 30°C, pH 7.2, 1000 rpm, and 30% dissolved oxygen. Four hours post-inoculation 50% glucose solution was fed into the bioreactor at 0.28 mL/min until an OD600 of 20 is achieved. The 2L culture was then chilled to 25°C until expression was induced with 1mL of 1M IPTG then incubated for an additional 18 hours at 25°C with agitation, after which the vessel was cooled to 15°C.

The cells were pelleted by centrifugation at 8000 RCF for 15 minutes. The supernatant was discarded, and the pellet was resuspended in resuspension/lysis buffer (50 mM Tris HCl, 100 mM NaCl, 10 mM MgCl_2_, 10 mM Imidazole, 5% Glycerol (v/v), pH 7.5). The resuspended pool was lysed via homogenization, and the resulting lysate was adjusted using IEX B buffer (50 mM Tris HCl, 2M NaCl, 10 mM MgCl_2_, 5% Glycerol (v/v), pH 7.5) to achieve a pool conductivity of approximately 30 mS/cm to ensure nuclease protein stability. The adjusted lysate was subsequently clarified by centrifugation at 15k rpm for 30 minutes at 4°C, and the supernatant was loaded onto pre-flushed and pre-equilibrated 120ZB08A Zeta Plus depth filters as an additional clarification step followed by membrane filtration using a Mustang E membrane filter. Column purification began with capture IMAC (cIMAC) by loading the depth filtered lysate onto a HisPur Ni-NTA column pre-equilibrated with IMAC A buffer (50 mM Tris HCl, 300 mM NaCl, 10 mM MgCl_2_, 10 mM Imidazole, 5% Glycerol (v/v), pH 7.5). The column was re-equilibrated with IMAC A buffer and washed with IMAC B buffer (50 mM Tris HCl, 300 mM NaCl, 10 mM MgCl_2_, 10 mM Imidazole, 5% Glycerol (v/v), 1% Triton (v/v), pH 7.5), then re-equilibrated with IMAC A buffer prior to elution. The protein was collected via isocratic elution using elevated imidazole buffer with IMAC D buffer (50 mM Tris HCl, 300 mM NaCl, 10 mM MgCl_2_, 250 mM Imidazole, 5% Glycerol (v/v), pH 7.5) and then spiked with TEV protease before incubating at 4°C for a minimum of 48 hours. The proteolyzed pool was diafiltered into IMAC A buffer via tangential flow filtration 1 (TFF1) using a 30 kDa MWCO hollow fiber filter to prepare the sample for isolation from the cleaved MBP tag. Column purification was continued with subtractive IMAC (subIMAC) by loading the TFF1 pool onto another HisPur Ni-NTA column pre-equilibrated with IMAC A buffer in a flowthrough mode. The resulting subIMAC pool was diluted to achieve a salt concentration of 1MAE column pre-equilibrated with IEX A buffer (50 mM Tris HCl, 150 mM NaCl, 10 mM MgCl_2_, 5% Glycerol (v/v), pH 7.5) in a flowthrough mode before being spiked with IEX B buffer to reach a salt concentration of 300 mM NaCl in preparation for the final column step. The adjusted load sample was then loaded via cation exchange chromatography (CEX) onto a Poros XS column pre-equilibrated with CEX A buffer (50 mM Tris HCl, 300 mM NaCl, 10 mM MgCl_2_, 5% Glycerol (v/v), pH 7.5). The column was re-equilibrated following sample loading and washed with elevated salt using CEX B buffer (50 mM Tris HCl, 500 mM NaCl, 10 mM MgCl_2_, 5% Glycerol (v/v), pH 7.5). The purified protein was collected via isocratic elution with high salt using CEX C buffer (50 mM Tris HCl, 750 mM NaCl, 10 mM MgCl_2_, 5% Glycerol (v/v), pH 7.5). The resulting CEX pool was then ultrafiltered and diafiltered into Final Formulation Buffer (20 mM Tris, 300 mM NaCl, 10 mM MgCl_2_, 1 mM TCEP, 5% Glycerol (v/v), pH 7.5) at a target final concentration of 10 mg/mL and stored at ≤ -60°C.

#### Purification of AsCas12a

His-MBP-AsCas12a was purified as described previously.^30^ The plasmid was transformed into BL21 (DE3) cells (New England Biolabs). A starter culture was grown overnight in Lysogeny Broth (LB) media containing 50 µg/mL kanamycin at 37°C and 250 rpm. The overnight culture was diluted 1:100 into 2L of LB media supplemented with antibiotic and was grown at 37°C and 225 rpm to an OD600 of ∼0.6. The culture was then chilled until expression was induced with 1 mM IPTG for 20 hours at 16°C and 180 rpm.Following induction, cells were pelleted and resuspended in Equilibration Buffer (20 mM HEPES pH 7.5, 1M NaCl, 0.5 mM TCEP, 5% glycerol) supplemented with 0.1% Tween-20 and cOmplete Protease Inhibitor Cocktail tablet (Roche). Cells were lysed via sonication (3s on, 3s off, 6 min, for 5 cycles) on ice. Immediately after lysis, 10 mM MgCl_2_ and 1X DNase I grade II (Roche) were added to the lysate, followed by incubation with stirring at 4°C for 30 minutes.Lysate was clarified by centrifugation at 18k rpm for 30 minutes at 4°C, and the supernatant was loaded onto a pre-equilibrated HisTrap HP Column (Cytiva). The column was washed with 5% Ni Elution Buffer (20 mM HEPES pH 7.5, 1M NaCl, 250 mM Imidazole, 5% glycerol) before elution using a linear gradient of Ni Elution Buffer. Protein-containing fractions were pooled and dialyzed with TEV protease in Dialysis Buffer (20 mM HEPES pH 7, 150 mM NaCl, 1 mM TCEP, 5% glycerol) overnight at 4°C to remove the MBP tag. The AsCas12a sample was then loaded onto a HiTrap SP HP column (Cytiva) that was pre-equilibrated with Low Salt Buffer (20 mM HEPES pH 7, 150 mM NaCl, 1 mM TCEP, 10% glycerol). The protein was eluted with a linear gradient of High Salt Buffer (20 mM HEPES pH 7, 1M NaCl, 0.5 mM TCEP, 10% glycerol), and fractions containing AsCas12a were pooled. The AsCas12a sample was then fractionated over a S200 Increase 10/300 GL column (Cytiva) pre-equilibrated with Low Salt Buffer supplemented with 5 mM MgCl_2_. Fractions were confirmed to contain Cas12a utilizing SDS-PAGE and pooled before each subsequent step or freezing. AsCas12a sample was concentrated to ∼89 µM, aliquoted, flash frozen with liquid nitrogen, and stored at -80°C.

### Nucleic Acid Preparation

DNA oligonucleotides and crRNAs were synthesized by Integrated DNA Technologies. DNA duplexes were formed by incubating target and non-target strand oligonucleotides in duplex buffer (10 mM Tris-HCl pH 7.5, 20 mM NaCl) at 90°C for 3 minutes, followed by slowly cooling to room temperature. When forming duplexes, the fluorescently labeled strand was mixed with the unlabeled strand at a 1:1.5 molar ratio. The crRNA was annealed according to the same protocol as the DNA duplexes in the same buffer. Detailed sequence information can be found in Supplementary Table S1.

### Cryo-EM Grid Preparation, Data Collection, and Processing

Cas12a-MG29-1 grids were prepared by mixing the pre-formed RNP with duplexed target at a 1:1.2 molar ratio in Cas12a-MG29-1 Cryo Buffer (20 mM Tris pH 7.5, 300 mM NaCl, 10 mM MgCl_2_) and incubating the sample at 37°C for 15 minutes. The sample was frozen at a final effector concentration of 10 ^µ^M. Carbon grids (Quantifoil 1.2/1.3) were glow-discharged for 30 seconds, and 3 uL of the sample was applied to the grid. By utilizing an FEI Vitrobot Mark IV at 4°C and 100% humidity, grids were blotted for 7 seconds with a blot force of 0 and plunge-frozen in liquid ethane.

The dataset was collected on an TFS Glacios (200 kV) cryo-electron microscope equipped with a Falcon 4 detector (Gatan) with a pixel size of 0.94 Å. All images were recorded with Serial EM.^35^ Movies were recorded with a total exposure time of 15.425 sec which resulted in an accumulated dosage of 49 e/Å^2^ split into 60 electron event representation (EER) fractions. Motion correction, contrast transfer function (CTF) estimation, and particle picking were performed with CryoSPARC Live (v4.3).^36^ All further data processing was performed in CryoSPARC (v4.3). A total of 1,689 movies were collected for the dataset.

Particles were initially picked utilizing blob picker, a minimum particle diameter of 100 Å, a maximum particle diameter set to 180 Å, and a minimum separation distance of 0.5 diameters. Particle picks were manually inspected to limit particle outliers. Final workflow is explained below and shown in Supplementary Figure S1. 717,423 particles were extracted with a box size of 320 pixels and Fourier cropped to 128 pixels. These particles were classified into 50 2D classes with a maximum resolution of 8 Å and a circular mask with a 150 Å diameter. Classification was performed with a batchsize per class of 500, 40 online-EM iterations, and force max over poses/shifts turned off. All other settings were default. After manual 2D class selection was performed, the remaining particles numbered 473,424 and were then utilized for ab-initio volume reconstruction and heterogenous refinement with 3 classes. Classes were manually inspected for resemblance to the ternary complex, and a single class of 154,706 particles was then re-extracted with a box size of 320 pixels and no Fourier cropping for use in NU (non-uniform refinement). The particles and mask from the NU job were tested for particle heterogeneity using 3D Variability with resolution filtered to 5 Å and displayed in cluster mode with 10 clusters. Clusters were manually inspected for REC2 domain docking, combined, and then refined with NU. The particle stack chosen for the final reconstruction contained 37,362 particles and resulted in a final ternary map with an overall resolution of 3.15 Å. Neither Global CTF Refinement nor Local CTF Refinement improved map quality. Further post-processing of the final sharpened map from CryoSPARC (v4.3)^36^ was performed with EMReady.^37^

### Model Building

An initial model of Cas12a-MG29-1 was folded by AlphaFold2^38^ and fit into the ternary map. Domains that did not initially fit well into the map were removed and replaced using rigid body fitting in ChimeraX (v1.9).^39^ The R-loop from a previously published AsCas12a 20-bp structure (8SFO)^30^ was also fit into the ternary map within ChimeraX (v1.9) and used as an initial model for the R-loop. The online Namdinator^40^ server was then used to further fit the model into the cryo-EM map. After Namdinator fitting, the structure was refined using a workflow wherein nucleic acid alterations and local fitting were performed in COOT (v0.9.8),^41^ modeling and further fit optimization was done in ISOLDE (v1.9),^42^ and iterative real-space refinement was performed in Phenix (v1.20.1).^43^ Structural figures were generated using ChimeraX (v1.9).^39^

### *In vivo* GFP Interference Assay

The *E. coli* based GFP interference was performed as in previous studies.^44,45^ Briefly, Cas12a nucleases were cloned into a plasmid containing a pBAD promoter, a CloDF13 origin of replication, and a chloramphenicol antibiotic resistance gene. The crRNA was cloned into an orthogonal plasmid containing a ColE1 origin of replication under the control of a pBAD promoter and a carbenicillin antibiotic resistance gene. A third plasmid expressing sf-GFP was also cloned into a p15a low copy number plasmid with a native tac promoter for constitutive expression and a streptomycin resistance gene. Dh5^α^ (New England Biolabs) cells were transformed with the sf-GFP plasmid and the appropriate crRNA plasmid. Transformed cells were plated and selected for using LB plates containing chloramphenicol and streptomycin. After overnight growth, colonies were selected from the double antibiotic plates and grown up in the appropriate antibiotics to make the cells competent using the Mix & Go! E. coli transformation Kit (Zymo). Cells were then finally chemically transformed with the Cas12a nuclease plasmid and plated on LB plates containing chloramphenicol, streptomycin, and carbenicillin.

Three separate cultures were grown for 16-18 hours at 37°C and 250 rpm in chloramphenicol, carbenicillin, and streptomycin. Cultures were diluted to an OD600 of ∼0.05 in media containing chloramphenicol and carbenicillin. Additionally, 0.2% arabinose was added to half of the cultures to compare induced samples to uninduced. 350 ^µ^L of the diluted culture was plated in a black-walled, clear bottom 96-well plate (Invitrogen) and allowed to grow while shaking at 37°C for 17.5 hours in a BioTek Cytation 5 microplate reader (Agilent). Every 5 minutes, fluorescence emission at 515 nm after excitation at 470 nm and absorbance at 600 nm was monitored. To compare samples, endpoint GFP fluorescence normalized to cell density (GFP/OD) values were calculated for uninduced and induced cultures. Fold depletion (FD) was calculated as:

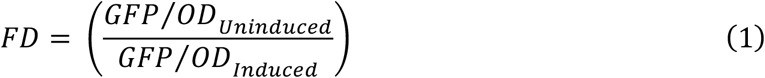

To quantify specificity of a certain mutant, a specificity score (SS) was calculated for each biological replicate as the difference between fold depletion values obtained for the perfectly matched target (PT) and the mismatched target (MM):

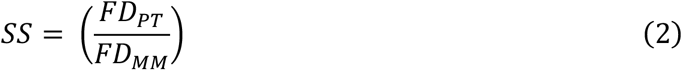

For statistical analyses, log_2_(FD) or log_2_(SS) values were calculated for each biological replicate and genotype. In comparisons involving more than two genotypes, one-way ANOVA followed by Dunnett’s multiple comparisons test (using WT Cas12a-MG29-1 as the control) was performed. For comparisons between two genotypes (AsCas12a and the loop mutant), unpaired t-tests with Welch’s correction were used. Data are presented as mean ± SEM from three independent biological replicates.

### Measurement of DNA Cleavage Rates

For substrate-initiated cleavage reactions, RNP complexes were formed by incubating Cas12a-MG29-1 with a 1.5x molar excess of crRNA in Magnesium Reaction Buffer (20 mM Tris pH 7.5, 150 mM NaCl, 5% glycerol, 1 mM TCEP, and 10 mM MgCl_2_) for 15 min at room temperature. Assembled RNP complexes were further diluted in Magnesium Reaction Buffer and pre-incubated at 37°C for 30 minutes. To initiate the reactions, 5’FAM-labeled target duplex was mixed with active protein to final concentrations of 10 nM and 50 nM, respectively. Reactions were quenched at various timepoints with 0.5 M EDTA.

For pre-formed R-loop cleavage reactions, RNP complexes were formed as in substrate-initiated reactions with the exception that the assembly occurred in Reaction Buffer without magnesium (20 mM Tris pH 7.5, 150 mM NaCl, 5% glycerol, and 1 mM TCEP) supplemented with 0.3 mM EDTA. Assembled RNP complexes were then diluted in Reaction Buffer without magnesium and incubated with 10 nM 5’FAM-labeled target duplex at 37°C for 30 min. These reactions were initiated using Mg^2+^, with final reaction concentrations of 10.3 mM Mg^2+^ and 50 nM active Cas12a-MG29-1 protein. Reactions were quenched as described for substrate-initiated cleavage. AsCas12a cleavage reactions were performed similarly with the exception that the cleavage reaction buffers were supplemented with 0.2 mg/mL BSA. All above cleavage reactions were performed in a 37°C water bath or heating block. Cleavage products were resolved via capillary electrophoresis using an Applied Biosystems DNA sequencer (ABI 3130xI) with POP-7. Traces corresponding to uncleaved substrate and product were quantified with GeneMapper software (v6) to determine amount of product formed over time. Each reaction was repeated at least twice.

### Measurement of R-loop Propagation Rates

R-loop propagation rates were measured with a stopped-flow assay as previously described for Cas9.^46^ Substrate DNA oligonucleotides containing the tricyclic fluorescent cytidine analog (tC°) were synthesized and PAGE purified by Bio-Synthesis, Inc. These substrates were combined with the corresponding unlabeled strand and duplexed according to the protocol described in nucleic acid preparation above. 100 nM Cas12a-MG29-1 or AsCas12a RNP complex was mixed with 50 nM tC°-labeled target duplex at 37°C in an AutoSF-120 stopped-flow instrument (KinTek Corporation, Austin, TX). Lamp excitation was at 367 nm, and emission was measured using a 445 nm filter with a 20 nm bandpass filter (Semrock). Measurements were repeated 2 times.

### Global fitting of kinetic data

Kinetic data global fitting was performed by fitting all experiments in KinTek Explorer^47^ (KinTek Corporation, Austin, TX) using the reaction scheme in Figure 4B. Each experiment was modeled exactly as it was performed by inputting mixing steps, initial reactant concentrations, and defining output observables. Products of chemical quench experiments were modeled as the sum of species containing product that was labeled in the experiment (i.e. for NTS cleavage: EDP1+EDP2). The stopped flow fluorescence data measuring R loop formation/cleavage was modeled using fluorescence scaling factors, as shown below, where *a* scales the overall signal, *b1* represents the fractional change in fluorescence in forming the R-loop completed state, and *b2* represents the fractional change in fluorescence in forming the cleaved product state:

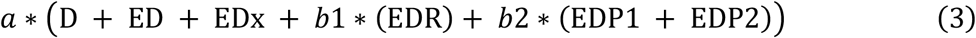

For the DNA binding step, DNA on and off rates were not well constrained by the data so the binding rate constant was locked at 500 ^µ^M^-1^ s^-1^ as to not limit the rate of R-loop formation and the reverse rate constant as locked at 0.01 s^-1^, giving a tight K_d_, as estimated in previous studies.^29^

Confidence contours shown in Supplementary Figure S2 were calculated using the FitSpace function in KinTek Explorer. These confidence contour plots are calculated by systematically varying a single rate constant and holding it fixed at a particular value while refitting the data, allowing all other rate constants to float. The goodness of fit was scored by the resultingχ^2^ value. The confidence interval is defined based on a threshold in χ^2^ calculated from the F-distribution based on the number of data points and number of variable parameters to give the 95% confidence limits.^48^ For the data in Figure 3, this threshold was 0.99 and was used to estimate the upper and lower limits for each rate constant which are reported in Supplementary Table S2.

The free energy profile in Figure 4 was created in KinTek Explorer using the rate constants given in Supplementary Table 2 using simple transition state theory:

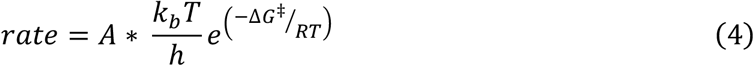

where *k*_*b*_ is the Boltzmann constant, *h* is Planck’s constant, T is temperature, and R is the gas constant. The free energy profile was created for the reaction at 37°C using a transmission coefficient of A=0.01 and a value for DNA concentration of 1 nM for the second order DNA binding step.

## RESULTS

### Structure of the Cas12a-MG29-1 ternary complex

To investigate the structural basis for the enhanced activity and specificity of Cas12a-MG29-1, we determined the cryo-EM structure of Cas12a-MG29-1 bound to a crRNA and its DNA target containing a T-rich PAM. The structure captures the product state of the Cas12a-MG29-1-crRNA-target complex at a nominal resolution of ∼3.2Å (Fig. 1B-1D, Supplementary Fig. S1, and Supplementary Table S3). Overall, our model of Cas12a-MG29-1 adopts the bilobed architecture typical of other Cas12a orthologs, consisting of the recognition (Rec) and nuclease (NUC) lobes. The Rec lobe contains the REC1 and REC2 domains, which flank the crRNA-DNA heteroduplex as it traverses the cleft between the two lobes. The NUC lobe includes the PAM-interacting (PI), wedge (WED), bridge helix (BH), RuvC, and Nuc domains, which accommodate the path of the displaced non-target strand (NTS) as it is threaded toward the nuclease active site. The RuvC domain, together with the adjacent Nuc domain, forms the catalytic core of the enzyme, while the PI, WED, and BH domains form part of the structural scaffold that surrounds the R-loop and adjacent duplex DNA. In the Cas12a-MG29-1 structure, the catalytic center is positioned at the junction of the Nuc and RuvC domains, as in other Cas12a orthologs. Conserved acidic residues within the RuvC domain (D886, E976, and D1229) define the core of the nuclease active site (Supplementary Fig. S3A and S3B).

Given the central role of protein-nucleic acid interactions in governing Cas12a activity and specificity, we used the Cas12a-MG29-1 ternary structure to map contacts between the enzyme and the crRNA-DNA heteroduplex along the length of the R-loop (Fig. 1E). Because Cas12a nuclease activation occurs only after progression through late stages of R-loop propagation,^29,30,32,49^ contacts formed along the heteroduplex provide a structural reference for features that may influence target discrimination. A flexible loop (residues 259-271) adjacent to the heteroduplex at positions 14-16, hereafter referred to as Loop 1, is present in the structure but forms relatively limited contacts with the R-loop in this region (Fig. 1E).^30^ This is consistent with reduced stabilization during intermediate stages of R-loop formation that may support a stringent target search. In contrast, a second loop (Loop 2, residues 511-524) contacts the distal portion of the R-loop near positions 16-17, conferring stabilizing interactions near the late-forming region of the heteroduplex (Fig. 1E). Together, these features illustrate how flexible loop regions of Cas12a-MG29-1 are positioned relative to the R-loop and provide a structural reference for subsequent comparative analysis.

### Distinct R-loop architecture near the distal R-loop

To visualize the structural basis for the differences in R-loop contacts, we superimposed Cas12a-MG29-1 with the well-characterized Cas12a ortholog, AsCas12a.^30^ The overall architectures are highly similar (Fig. 2A); however, significant deviations emerge near the distal end of the R-loop. In this region, Loop 1 of Cas12a-MG29-1 is shifted away from the heteroduplex relative to AsCas12a, consistent with the reduced interactions at positions 14-16. Loop 2 also adopts a conformation that approaches the distal R-loop and forms contacts that are absent in AsCas12a (Fig. 2B and Supplementary Fig. S4A). Notably, AsCas12a does not contain a comparably positioned loop that extends toward the R-loop, effectively lacking this interaction surface. A structurally analogous Loop 2 is present in FnCas12a but although sufficiently extended to approach the distal duplex does not form comparable contacts, further highlighting variability in this region (Supplementary Fig. S4B and S4C).^32^ These differences in R-loop contacts are localized to flexible loop regions and occur without substantial rearrangement of the surrounding domains, highlighting structural variability near the late-forming portion of the heteroduplex.

**Figure 2.**
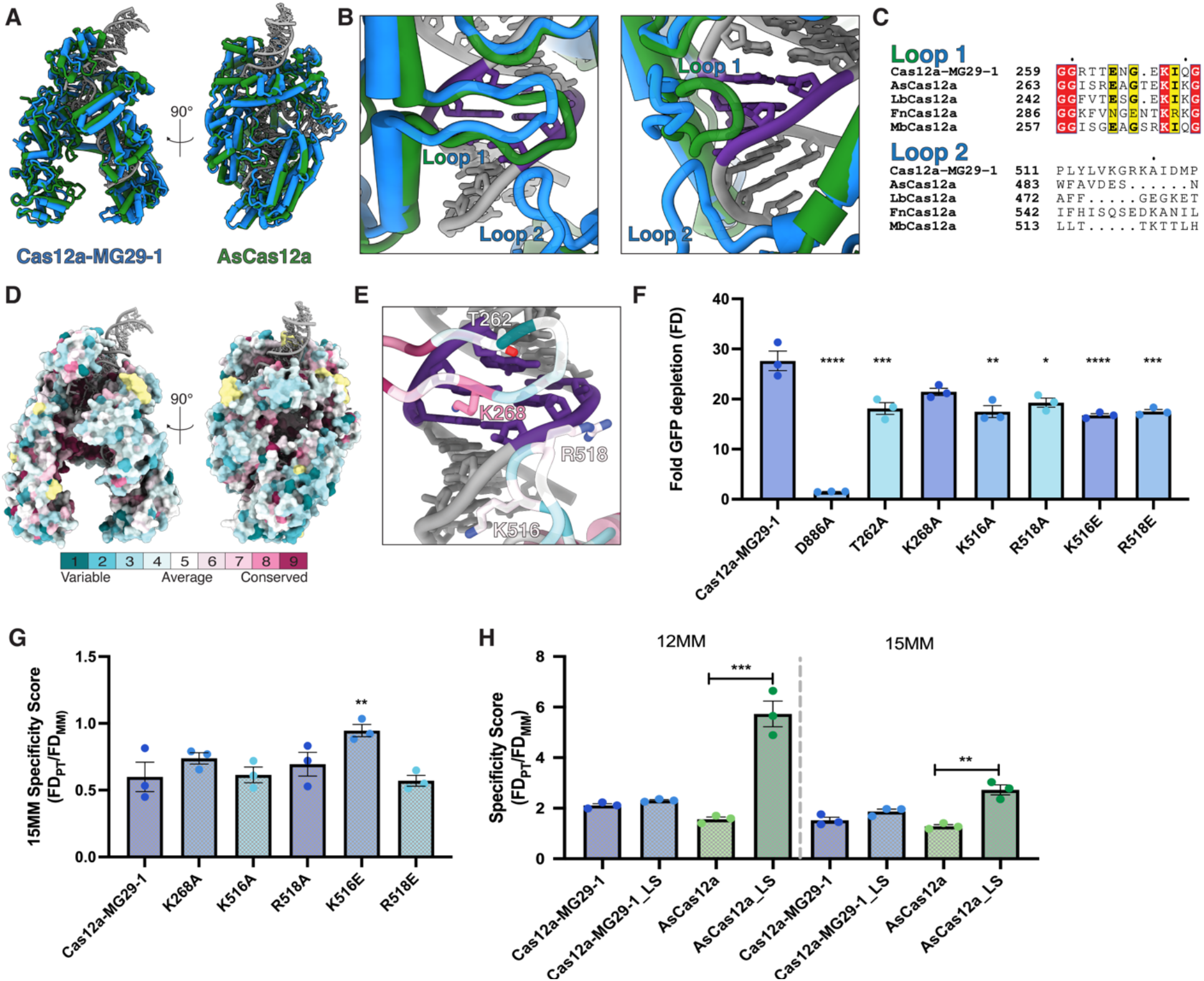
Variable distal R-loop loop architecture influences Cas12a activity and target discrimination. (A) Structural overlays of the Cas12a-MG29-1 model with AsCas12a (8SFR).^30^ Cas12a-MG29-1 is shown in blue, while AsCas12a is shown in green. The R-loop is colored gray. Front and side views are shown. (B) Close up view of Loop 1 and Loop 2. Positions 14-16 of the heteroduplex are highlighted in purple, while the rest of the figure matches the color scheme in (A). (C) Conservation of Loop 1 and Loop 2 in representative Cas12a orthologs.Highly conserved residues are highlighted in red, while yellow denotes partial conservation. Generated with ESPript.^50^ (D) Front and side views of the Cas12a-MG29-1 model surface representation colored by conservation score. Higher conservation is denoted by pink, while more variable positions are colored in blue. (E) Close-up view of Loop 1 and Loop 2 colored by conservation score. Residues chosen for mutagenesis studies are labeled. (F) GFP depletion assay with perfect target. One-way ANOVA with Dunnet’s multiple comparisons test;^****^p<0.0001, ^***^0.0001>p<0.001, ^**^0.001>p<0.01, ^*^0.01>p<0.05. All experiments were performed in biological triplicate. (G) GFP depletion assay with target plasmid containing target containing a mismatch (MM) at position 15. Statistics performed as in (F); and experiments were performed in biological triplicate. (H) GFP depletion assay with loop swap mutants. For Cas12a-MG29-1 and the loop swap mutant (Cas12a-MG29-1_LS), statistical tests were performed as in (F). For AsCas12a and the loop swap mutant (AsCas12a_LS), unpaired t-tests with Welch’s correction were performed. Statistical significance is denoted as in (F).

Sequence alignments of Loops 1 and 2 across representative Cas12a orthologs, including AsCas12a, FnCas12a, LbCas12a, and MbCas12a, revealed limited conservation, particularly within Loop 2 (Fig. 2C). Within Loop 1, only a single lysine residue (K268 in Cas12a-MG29-1) is conserved across all five orthologs. Additional positions show conservation of glutamic acid (264), glycine (259,260,271), or isoleucine residues (269); however, these amino acids are less consistent with direct interactions with the nucleic acid backbone. Loop 2 varies in both length and composition across orthologs, consistent with structural variability observed in this region.

To further evaluate conservation, we expanded the search to the ConSurf server,^51^ which identified 101 structural orthologs to Cas12a-MG29-1 and mapped sequence conservation across the entire sequence of the protein. Both loops displayed low overall conservation, with only K268 exhibiting high conservation. The variability of these regions suggests that distal R-loop contacts do not depend on strict amino acid sequence identity but instead arise from flexible structural elements that may contribute to target recognition. These observations guided subsequent mutagenesis experiments designed to test whether the altered loop engagement in Cas12a-MG29-1 contributes to functional differences in activity and/or target discrimination.

### Functional analysis of loop-mediated contacts at the distal R-loop

To test whether Loop 1 or Loop 2 residues contribute to Cas12a-MG29-1 activity and specificity, we performed structure-guided mutagenesis of key residues within these loops. Despite limited conservation (Fig. 2E), K268 and T262 in Loop 1 and K516 and R518 in Loop 2 were selected based on proximity to the R-loop. Alanine substitutions were generated at each position, along with charge-reversal mutants K516E and R518E, and evaluated using a GFP depletion assay (Fig. 2F). In this assay, Cas12a-mediated cleavage of an sfGFP plasmid reduces fluorescent signal, such that a lower GFP fluorescence corresponds to higher nuclease activity. A catalytically dead mutant (D886A) showed minimal fold GFP depletion. In contrast, the alanine and charge-reversal mutants reduced GFP depletion relative to wild type, except K286A, which did not show a significant effect, indicating that residues within both loops contribute to nuclease activity.

To assess mismatch tolerance, these variants were tested against a target containing a mismatch at position 15 and compared to the perfect target in the GFP depletion assay. Specificity was evaluated by comparing the relative depletion to the perfectly matched and mismatched targets, providing a measure of how effectively each variant discriminates against mismatched substrates in a cellular context. Most substitutions did not significantly alter specificity; however, K516E produced a significant increase in discrimination, suggesting that this contact contributes to mismatch surveillance within the distal R-loop.

To test whether specificity arises from loop architecture rather than individual residues, we performed Loop 1 swaps between Cas12a-MG29-1 and AsCas12a (Fig. 2H). Introducing the Cas12a-MG29-1 Loop 1 into AsCas12a (AsCas12a_LS) did not reduce activity on a perfect target (Supplementary Fig. S4D) but increased mismatch discrimination at both position 12 and position 15. In contrast, replacing Loop 1 of Cas12a-MG29-1 with that of AsCas12a (Cas12a-MG29-1_LS) did not measurably alter on-target activity or mismatch tolerance. These results indicate that the Cas12a-MG29-1 Loop 1 can enhance specificity in the AsCas12a background but is not sufficient on its own to account for the specificity of Cas12a-MG29-1, suggesting that distal R-loop surveillance is influenced by the broader structural context. This context dependence, together with the effect of Loop 2 substitutions, prompted us to investigate whether these differences alter the kinetics of R-loop formation and cleavage.

### Kinetic comparison of Cas12a-MG29-1 and AsCas12a

To further characterize the differences between Cas12a-MG29-1 and AsCas12a targeting, we performed single-turnover kinetic analysis using in vitro cleavage assays and stopped-flow measurements of R-loop formation (Fig. 3). Cleavage kinetics were monitored using DNA substrates labeled with 5’-6-FAM (Fluorescein) on either the NTS or target strand (TS).Reactions were performed under two conditions. In the first, cleavage was initiated by mixing pre-formed ribonucleoprotein (RNP) complexes with target DNA, such that the DNA binding and R-loop formation steps must precede cleavage. In the second, RNP and DNA were preincubated in the absence of Mg^2+^ to allow formation of a cleavage-competent complex, and reactions were initiated by addition of Mg^2+^.

**Figure 3.**
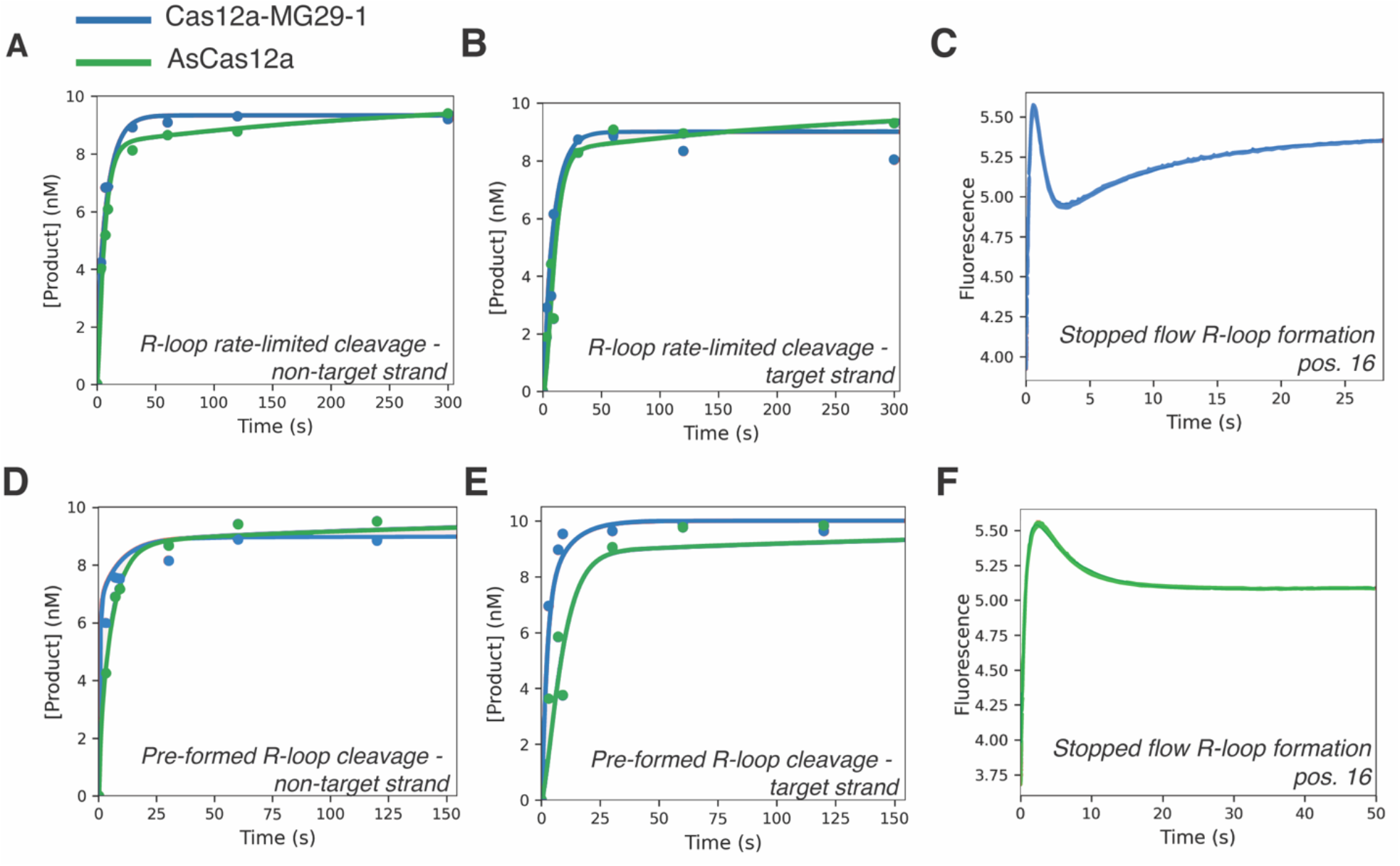
Cas12a-MG29-1 displays faster DNA cleavage than AsCas12a with multiphasic R-loop formation. In all panels, Cas12a-MG29-1 data are colored in blue, while AsCas12a data are colored in green. (A) DNA-initiated cleavage of the non-target strand of DNA. Cas12a ribonucleoprotein (RNP) was mixed with a DNA target containing fluorescent label on the non-target strand in the presence of Mg^2+^ to start the reaction. (B) DNA-initiated cleavage of the target strand of DNA. Reaction conditions were the same as in (A) except the fluorescent label was on the target strand of the DNA duplex. (C) R-loop formation of Cas12a-MG29-1. R-loop formation was initiated by mixing a DNA duplex with a tC° base at position 16 with Cas12a-MG29-1, and the change in fluorescence was measured via stopped-flow. (D) Mg^2+^-initiated cleavage of the non-target strand DNA. Cas12a RNP and DNA were mixed with Mg^2+^ to start the reaction. The non-target strand was labeled. (E) Mg^2+^-initiated cleavage of the target strand DNA. Reaction conditions were the same as in (D), except the target strand was labeled. (F) R-loop formation of AsCas12a. R-loop formation was initiated by mixing a DNA duplex with a tC° base at position 16 with AsCas12a, and the change in fluorescence was measured via stopped-flow. For (A-F), data points represent a single representative dataset used for global data fitting, and the lines through the data represent the best fit obtained using KinTek Explorer software.^47^ Experiments were performed in duplicate to confirm reproducibility. For stopped-flow measurements (C,F), data were collected at high temporal resolution, such that individual data points are not visually resolved at the scale shown. The model and corresponding rates are summarized in Fig. 4.

We first examined DNA-initiated cleavage, and under these conditions Cas12a-MG29-1 cleavage was monophasic, while AsCas12a displayed biphasic behavior (Fig. 3A and B). Cas12a-MG29-1 product formation increased rapidly and approached a plateau in a single apparent phase, whereas AsCas12a showed an initial rapid increase followed by slower continued product accumulation, consistent with biphasic cleavage behavior. The multiphasic nature of these traces complicates equation-based fitting and direct comparison between the two orthologs since the observed rates are composites of multiple rate constants. To directly evaluate cleavage kinetics independent of preceding steps, we next measured cleavage after pre-forming the R-loop and initiating reactions with Mg^2+^(Fig. 3D and E). For both enzymes, Mg^2+^-initiated cleavage proceeded more rapidly than DNA-initiated cleavage, indicating that a step preceding chemistry is rate-limiting under these conditions.^29^ Under both reaction conditions, Cas12a-MG29-1 cleaved DNA more rapidly than AsCas12a.

We next measured the rate of R-loop formation using a stopped-flow fluorescence assay previously developed for SpCas9 and subsequently applied to multiple CRISPR–Cas systems (Fig. 3C and F).^46,52,44,53^ The fluorescent nucleoside analog tC° (1,3-diaza-2-oxophenoxazine) was incorporated at position 16 in the NTS of the DNA substrate such that R-loop formation causes the fluorophore to transition from a double-stranded to single-stranded environment, producing a fluorescence change. Unlike previous measurements for SpCas9, which show an increase in fluorescence during R-loop formation, both Cas12a-MG29-1 and AsCas12a exhibited an initial increase in fluorescence followed by a smaller decrease, consistent with additional conformational or cleavage-dependent steps following R-loop formation. In addition, both cleavage and R-loop formation traces displayed multiphasic behavior, indicating contributions from multiple steps in the reaction pathway. Because these data represent composite contributions from R-loop formation, conformational rearrangements, and cleavage, the observed rates are difficult to interpret using equation-based fitting. To more clearly define the underlying kinetic steps and compare the two enzymes, we globally fit the cleavage and stopped-flow fluorescence data for each enzyme with a simulation-based fitting approach (Fig. 4).

### Global kinetic modeling of Cas12a-MG29-1 and AsCas12a

We globally fit all kinetic data with KinTek Explorer software^47^ to a model that simultaneously accounts for DNA binding, R-loop formation, and sequential strand cleavage (Fig. 4). DNA binding steps were locked at identical values for both enzymes since these rate constants were not well constrained by the data. After DNA binding, the model incorporates reversible R-loop formation followed by sequential cleavage of the NTS and TS (Fig. 4B). This arrangement is consistent with the single RuvC nuclease active site in Cas12a and prior observations that cleavage of the NTS precedes TS cleavage.^29,32^ The biphasic cleavage behavior further required inclusion of an additional step, modeled as an ED to ED_x_ transition prior to cleavage, consistent with intermediates described for Cas9d and Cas12f.^44,53^ The biphasic stopped flow fluorescence traces were accounted for by one fluorescence scaling factor for the R loop formed state, EDR, and another scaling factor for the cleaved product states, EDP1 and EDP1P2. Scaling factors for these states were all higher than 1, indicating that they had higher fluorescence than the D, ED, and ED_x_ states, consistent with the double stranded to single stranded transition.

**Figure 4.**
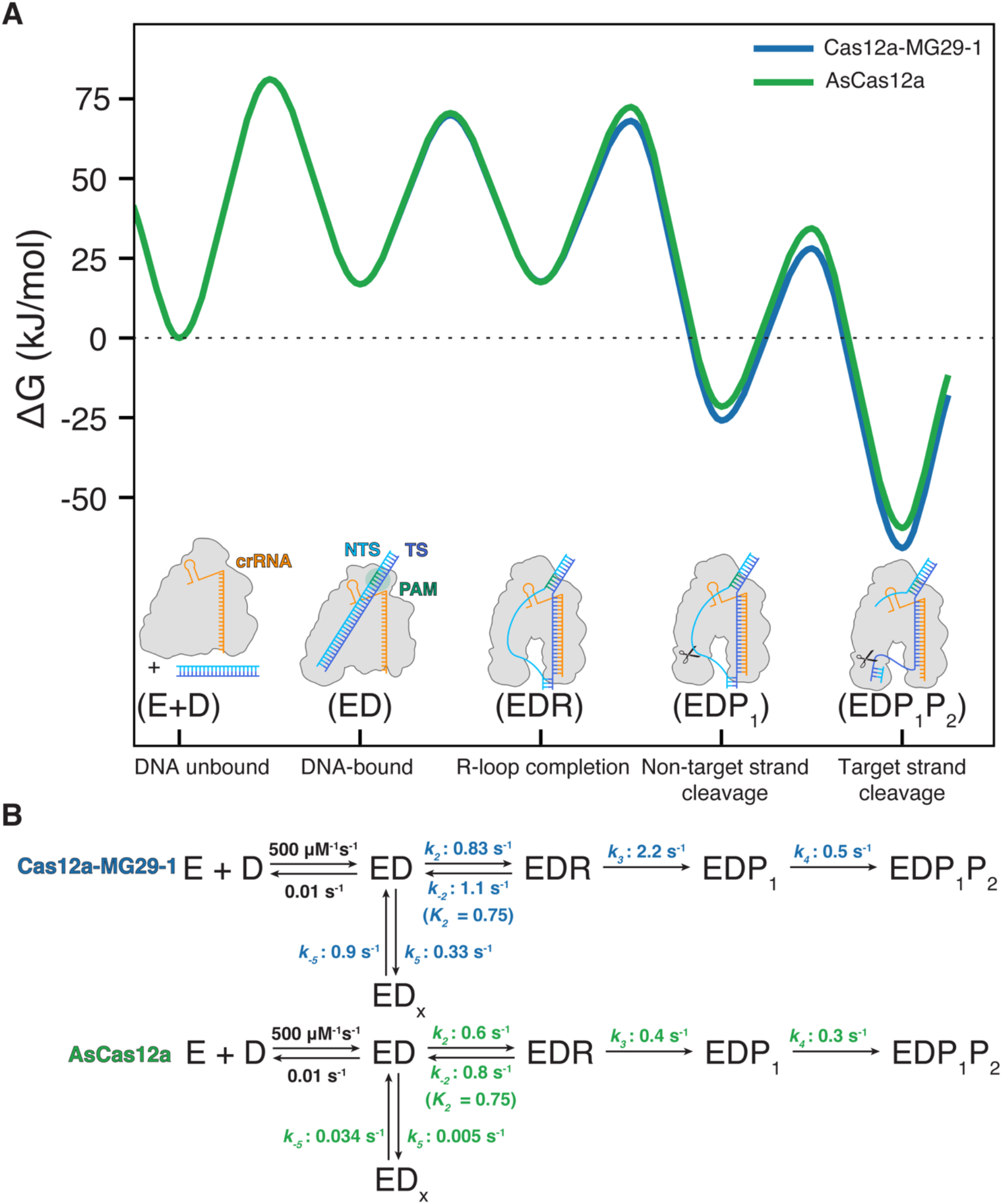
Kinetic partitioning after reversible R-loop formation distinguishes Cas12a-MG29-1 from AsCas12a. (A) Free energy profile of the Cas12a-MG29-1 and AsCas12a cleavage mechanisms. The Cas12a-MG29-1 model is shown in blue, whereas AsCas12a is shown in green. (B) Sequential kinetic model for Cas12a-MG29-1 and AsCas12a. Global kinetic fitting was performed as described with confidence contours provided in Supplementary Fig. S2.

Rate constants derived from global fitting gave valuable insights into mechanistic differences between the enzymes. Surprisingly, similar R-loop formation kinetics were obtained for Cas12a-MG29-1 and AsCas12a. Both enzymes exhibited forward R-loop formation rates of approximately 0.6–0.83 s^−1^ and reverse rates between 0.8 and 1.1 s^−1^, corresponding to comparable equilibrium constants of ∼0.75 for this step (Fig. 4B). These results indicate that both enzymes form similarly reversible R-loops.

In contrast, cleavage rates differed substantially between the two enzymes. Global kinetic modeling indicated that NTS cleavage occurred at 2.2 s^−1^ for Cas12a-MG29-1 compared to 0.4 s^−1^ for AsCas12a, representing the largest divergence in the kinetic pathway (Fig. 4B and Supplementary Table S2). Subsequent TS cleavage proceeded at 0.5 s^−1^ for Cas12a-MG29-1 and 0.3 s^−1^ for AsCas12a. These fitted differences correspond to free energy profiles with similar barriers for R-loop formation but reduced barriers for strand cleavage in Cas12a-MG29-1 (Fig. 4A).

Both enzymes exhibit reversible R-loop formation, allowing dissociation prior to irreversible cleavage. Although R-loop forward and reverse rates are similar for both enzymes, cleavage rates differ, resulting in distinct kinetic partitioning of the EDR state. Cas12a-MG29-1 more efficiently partitions the R-loop intermediate toward product formation, with NTS cleavage occurring at 2.2 s^−1^, nearly twice the rate of R-loop reversal (1.1 s^−1^). This kinetic partitioning favors product formation and leads to efficient on-target cleavage. In contrast, AsCas12a shows slower cleavage at 0.4 s^−1^, while the rate of R-loop reversal is 0.8 s^−1^, indicating that on-target DNA has an opportunity to dissociate before cleavage, lowering cleavage efficiency. These differences in kinetic partitioning explain the increased cleavage efficiency observed for Cas12a-MG29-1 while maintaining reversible R-loop formation. This suggests that faster, irreversible commitment to cleavage may enhance target discrimination by promoting progression of stable R-loops while allowing less stable intermediates to dissociate prior to catalysis.

## DISCUSSION

Understanding how Cas12a enzymes balance efficient target cleavage with discrimination is important for both mechanistic insight and the development of improved genome editing technologies. Cas12a-MG29-1 exhibits enhanced activity and specificity relative to AsCas12a, yet the mechanistic basis for this behavior has remained unclear.^27,28^ In this study, structural, functional, and kinetic analyses revealed differences in distal R-loop interactions that influence catalytic commitment without substantially altering R-loop formation. The Cas12a-MG29-1 structure reveals repositioning of flexible loop elements near the late-forming region of the heteroduplex, including reduced engagement of Loop 1 and additional contacts formed by Loop 2. These regions display limited sequence conservation across Cas12a orthologs, suggesting that distal R-loop interactions may depend more on positioning and flexibility of the loops than on specific conserved amino acid identities. Mutational analysis of residues within these loops reduced activity without substantially altering mismatch discrimination, whereas transplantation of Loop 1 from Cas12a-MG29-1 into AsCas12a increased specificity without compromising activity. However, replacing Loop 1 of Cas12a-MG29-1 with that of AsCas12a did not reduce specificity, indicating that Loop 1 alone is not sufficient to account for the behavior of Cas12a-MG29-1. Together, these findings suggest that loop architecture near the distal R-loop modulates target discrimination in a context-dependent manner, likely through collective structural effects rather than individual conserved residues or a single loop element.

Kinetic analysis further revealed that Cas12a-MG29-1 and AsCas12a exhibit similar rates and reversibility of R-loop formation on perfect targets, indicating that initial target interrogation proceeds through comparable pathways. Instead, the primary kinetic distinction arises following R-loop formation, where Cas12a-MG29-1 displays substantially faster NTS cleavage. Because R-loop formation is reversible, cleavage competes with R-loop collapse, creating a kinetic partitioning step that determines whether a bound substrate proceeds to product formation or dissociates. The increased rate of NTS cleavage in Cas12a-MG29-1 shifts this partitioning toward productive cleavage while preserving reversible R-loop formation. In contrast, the slower cleavage rate of AsCas12a favors reversal of the R-loop intermediate, reducing overall activity.

Consistent with this interpretation, recent work has shown that subtle alterations in protein-DNA interactions can substantially enhance Cas12a catalytic efficiency without large changes in overall structure.^54^ A recent study of an engineered AsCas12a variant demonstrated that reducing steric constraints on the DNA can facilitate access to cleavage-competent conformations. In contrast to these findings, where increased activity arises from faster R-loop formation, our results indicate that Cas12a-MG29-1 achieves enhanced activity through accelerated cleavage, highlighting that distinct steps along the reaction pathway can be tuned to modulate Cas12a function.

These findings suggest that enhanced specificity in Cas12a-MG29-1 may arise not from improved discrimination during R-loop formation, but from accelerated irreversible commitment to cleavage. Both enzymes sample targets through reversible R-loop formation, allowing mismatched substrates to dissociate prior to cleavage. In principle, the faster cleavage rate of Cas12a-MG29-1 could increase the likelihood that stable R-loops proceed to cleavage before reversal, while less stable intermediates dissociate prior to catalysis. In this manner, specificity may be influenced through kinetic partitioning rather than increased binding discrimination.

This behavior is conceptually similar to nucleotide selection by DNA polymerases, where fidelity is achieved not only through base-pair recognition but also through accelerated chemistry following correct substrate binding.^55,56^ In polymerases, correctly paired nucleotides are incorporated more rapidly than mismatched substrates, shifting kinetic partitioning toward product formation. Although mismatch-dependent kinetics were not directly measured here, Cas12a-MG29-1 and AsCas12a exhibited similar R-loop formation kinetics, while Cas12a-MG29-1 displayed faster irreversible commitment to cleavage following formation of a stable R-loop. This comparison highlights how differences in post-R-loop kinetic steps could, in principle, contribute to differences in discrimination without requiring changes in initial target recognition.

Together, these findings support a model in which distal R-loop interactions modulate the transition from reversible target interrogation to irreversible cleavage. Rather than being governed by a single structural determinant, differences in loop architecture appear to influence this transition, while kinetic analysis indicates that enhanced activity arises from accelerated strand cleavage rather than substantial changes in R-loop formation kinetics. These observations raise the possibility that kinetic partitioning contributes to how Cas12a enzymes balance efficient target cleavage with discrimination against less stable R-loop intermediates. Together, these results provide a mechanistic framework for interpreting functional differences among Cas12a orthologs and inform the development of next-generation CRISPR enzymes.

## ACKNOWLEDGEMENTS

We thank Dr. Daniel Dickinson, the Dickinson and Taylor labs, and Metagenomi Therapeutics, Inc. for insightful discussions around the project. We also thank Andrew Langtry, Jacob Hogben, Neil Sanchez, and Daniel Chen from Metagenomi Therapeutics, Inc. for contributing to the protein expression and purification development of Cas12a-MG29-1. We thank Dr. Axel Brilot and Dr. Evan Schwartz at the Sauer Structural Biology Lab at UT Austin for assistance with cryo-EM. We thank Matt Hooper for development of the *in vivo* GFP assay.

## AUTHOR CONTRIBUTIONS

Jacquelyn T. Wright: Conceptualization, Data curation, Formal analysis, Investigation, Methodology, Writing—original draft, Writing—review & editing. Satya Yendluri: Methodology, Resources. Nicole C. Thomas: Project administration. Cristina N. Butterfield: Supervision, Writing—review & editing. Tyler L. Dangerfield: Formal analysis, Methodology, Writing—review & editing. David W. Taylor: Funding acquisition, Supervision, Writing— review & editing.

## SUPPLEMENTARY DATA

Supplementary Data are available online.

## DATA AVAILABILITY

The structures and associated atomic coordinates have been deposited into the Electron Microscopy Data Bank (EMDB) and the Protein Data Bank (PDB).

## REFERENCES

1. Barrangou, R. et al. CRISPR Provides Acquired Resistance Against Viruses in Prokaryotes. Science 315, 1709–1712 (2007).

2. Sorek, R., Lawrence, C. M. & Wiedenheft, B. CRISPR-Mediated Adaptive Immune Systems in Bacteria and Archaea. Annual Review of Biochemistry 82, 237–266 (2013).

3. van der Oost, J., Jore, M. M., Westra, E. R., Lundgren, M. & Brouns, S. J. J. CRISPR-based adaptive and heritable immunity in prokaryotes. Trends in Biochemical Sciences 34, 401–407 (2009).

4. Jinek, M. et al. RNA-programmed genome editing in human cells. eLife 2, e00471 (2013).

5. Mali, P. et al. RNA-Guided Human Genome Engineering via Cas9. Science 339, 823–826 (2013).

6. Wang, J. Y. & Doudna, J. A. CRISPR technology: A decade of genome editing is only the beginning. Science 379, eadd8643 (2023).

7. Koonin, E. V., Gootenberg, J. S. & Abudayyeh, O. O. Discovery of Diverse CRISPR-Cas Systems and Expansion of the Genome Engineering Toolbox. Biochemistry 62, 3465–3487 (2023).

8. Makarova, K. S. et al. An updated evolutionary classification of CRISPR–Cas systems including rare variants. Nat Microbiol 10, 3346–3361 (2025).

9. Fu, Y. et al. High-frequency off-target mutagenesis induced by CRISPR-Cas nucleases in human cells. Nat Biotechnol 31, 822–826 (2013).

10. Kleinstiver, B. P. et al. Engineered CRISPR-Cas9 nucleases with altered PAM specificities. Nature 523, 481–485 (2015).

11. Doudna, J. A. The promise and challenge of therapeutic genome editing. Nature 578, 229–236 (2020).

12. Pacesa, M., Pelea, O. & Jinek, M. Past, present, and future of CRISPR genome editing technologies. Cell 187, 1076–1100 (2024).

13. Kim, D. et al. Genome-wide analysis reveals specificities of Cpf1 endonucleases in human cells. Nat Biotechnol 34, 863–868 (2016).

14. Zetsche, B. et al. Cpf1 Is a Single RNA-Guided Endonuclease of a Class 2 CRISPR-Cas System. Cell 163, 759–771 (2015).

15. Swarts, D. C., van der Oost, J. & Jinek, M. Structural Basis for Guide RNA Processing and Seed-Dependent DNA Targeting by CRISPR-Cas12a. Molecular Cell 66, 221–233.e4 (2017).

16. Fonfara, I., Richter, H., Bratovič, M., Le Rhun, A. & Charpentier, E. The CRISPR-associated DNA-cleaving enzyme Cpf1 also processes precursor CRISPR RNA. Nature 532, 517–521 (2016).

17. Zetsche, B. et al. Multiplex gene editing by CRISPR–Cpf1 using a single crRNA array. Nat Biotechnol 35, 31–34 (2017).

18. Lee, J. G. et al. Knockout rat models mimicking human atherosclerosis created by Cpf1-mediated gene targeting. Sci Rep 9, 2628 (2019).

19. Hebert, J. D. et al. Efficient and multiplexed somatic genome editing with Cas12a mice. Nat.Biomed. Eng 9, 1982–1997 (2025).

20. Gier, R. A. et al. High-performance CRISPR-Cas12a genome editing for combinatorial genetic screening. Nat Commun 11, 3455 (2020).

21. Hsiung, C. C.-S. et al. Engineered CRISPR-Cas12a for higher-order combinatorial chromatin perturbations. Nat Biotechnol 43, 369–383 (2025).

22. Chen, J. S. et al. CRISPR-Cas12a target binding unleashes indiscriminate single-stranded DNase activity. Science 360, 436–439 (2018).

23. Li, S.-Y. et al. CRISPR-Cas12a has both cis- and trans-cleavage activities on single-stranded DNA. Cell Res 28, 491–493 (2018).

24. Rananaware, S. R. et al. Programmable RNA detection with CRISPR-Cas12a. Nat Commun 14, 5409 (2023).

25. Li, S.-Y. et al. CRISPR-Cas12a-assisted nucleic acid detection. Cell Discov 4, 20 (2018).

26. Broughton, J. P. et al. CRISPR–Cas12-based detection of SARS-CoV-2. Nat Biotechnol 38, 870–874 (2020).

27. Aliaga Goltsman, D. S. et al. Novel Type V-A CRISPR Effectors Are Active Nucleases with Expanded Targeting Capabilities. The CRISPR Journal 3, 454–461 (2020).

28. Lamothe, R. C. et al. Novel CRISPR-Associated Gene-Editing Systems Discovered in Metagenomic Samples Enable Efficient and Specific Genome Engineering. The CRISPR Journal 6, 243–260 (2023).

29. Strohkendl, I., Saifuddin, F. A., Rybarski, J. R., Finkelstein, I. J. & Russell, R. Kinetic Basis for DNA Target Specificity of CRISPR-Cas12a. Molecular Cell 71, 816–824.e3 (2018).

30. Strohkendl, I. et al. Cas12a domain flexibility guides R-loop formation and forces RuvC resetting. Molecular Cell 84, 2717–2731.e6 (2024).

31. Szczelkun, M. D. et al. Direct observation of R-loop formation by single RNA-guided Cas9 and Cascade effector complexes. Proceedings of the National Academy of Sciences 111, 9798–9803 (2014).

32. Stella, S. et al. Conformational Activation Promotes CRISPR-Cas12a Catalysis and Resetting of the Endonuclease Activity. Cell 175, 1856–1871.e21 (2018).

33. Gao, P., Yang, H., Rajashankar, K. R., Huang, Z. & Patel, D. J. Type V CRISPR-Cas Cpf1 endonuclease employs a unique mechanism for crRNA-mediated target DNA recognition. Cell Res 26, 901–913 (2016).

34. Yamano, T. et al. Crystal Structure of Cpf1 in Complex with Guide RNA and Target DNA. Cell 165, 949–962 (2016).

35. Mastronarde, D. N. Automated electron microscope tomography using robust prediction of specimen movements. Journal of Structural Biology 152, 36–51 (2005).

36. Punjani, A. Real-time cryo-EM structure determination. Microanal 27, 1156–1157 (2021).

37. He, J., Li, T. & Huang, S.-Y. Improvement of cryo-EM maps by simultaneous local and non-local deep learning. Nat Commun 14, 3217 (2023).

38. Jumper, J. et al. Highly accurate protein structure prediction with AlphaFold. Nature 596, 583–589 (2021).

39. Meng, E. C. et al. UCSF ChimeraX: Tools for structure building and analysis. Protein Science 32, e4792 (2023).

40. Kidmose, R. T. et al. Namdinator – automatic molecular dynamics flexible fitting of structural models into cryo-EM and crystallography experimental maps. IUCrJ 6, 526–531 (2019).

41. Emsley, P., Lohkamp, B., Scott, W. G. & Cowtan, K. Features and development of Coot. Acta Cryst D 66, 486–501 (2010).

42. Croll, T. I. ISOLDE: a physically realistic environment for model building into low-resolution electron-density maps. Acta Cryst D 74, 519–530 (2018).

43. Liebschner, D. et al. Macromolecular structure determination using X-rays, neutrons and electrons: recent developments in Phenix. Acta Cryst D 75, 861–877 (2019).

44. Ocampo, R. F. et al. DNA targeting by compact Cas9d and its resurrected ancestor. Nat Commun 16, 457 (2025).

45. Phan, P., Schelling, M., Xue, C. & Sashital, D. Fluorescence-based methods for measuring target interference by CRISPR–Cas systems. in Methods in Enzymology vol. 616 61–85 (Academic Press, 2019).

46. Liu, M.-S. et al. Engineered CRISPR/Cas9 enzymes improve discrimination by slowing DNA cleavage to allow release of off-target DNA. Nat Commun 11, 3576 (2020).

47. Global Kinetic Explorer: A new computer program for dynamic simulation and fitting of kinetic data. Analytical Biochemistry 387, 20–29 (2009).

48. Johnson, K. A., Simpson, Z. B. & Blom, T. FitSpace Explorer: An algorithm to evaluate multidimensional parameter space in fitting kinetic data. Analytical Biochemistry 387, 30–41 (2009).

49. Zhang, L. et al. Conformational Dynamics and Cleavage Sites of Cas12a Are Modulated by Complementarity between crRNA and DNA. iScience 19, 492–503 (2019).

50. Gouet, P. & Courcelle, E. ENDscript: a workflow to display sequence and structure information. Bioinformatics 18, 767–768 (2002).

51. Yariv, B. et al. Using evolutionary data to make sense of macromolecules with a “face-lifted” ConSurf. Protein Science 32, e4582 (2023).

52. Hibshman, G. N. et al. Unraveling the mechanisms of PAMless DNA interrogation by SpRY-Cas9. Nat Commun 15, 3663 (2024).

53. Guan, K. et al. Comparative characterization of Cas12f orthologs reveals mechanistic features underlying enhanced genome editing efficiency. 2025.08.14.670346 Preprint at 10.1101/2025.08.14.670346 (2025).

54. Jansson-Fritzberg, L. et al. Mechanistic basis for improved activity of Engineered AsCas12a. Commun Biol 10.1038/s42003-026-09799-1 (2026) doi:10.1038/s42003-026-09799-1.

55. Dangerfield, T. L. & Johnson, K. A. Conformational dynamics during high-fidelity DNA replication and translocation defined using a DNA polymerase with a fluorescent artificial amino acid. Journal of Biological Chemistry 296, (2021).

56. Dangerfield, T. L., Kirmizialtin, S. & Johnson, K. A. Conformational dynamics during misincorporation and mismatch extension defined using a DNA polymerase with a fluorescent artificial amino acid. Journal of Biological Chemistry 298, (2022).

