## Supplementary Information for "Structural and kinetic insights into a metagenomics-derived Cas12a with high specificity"

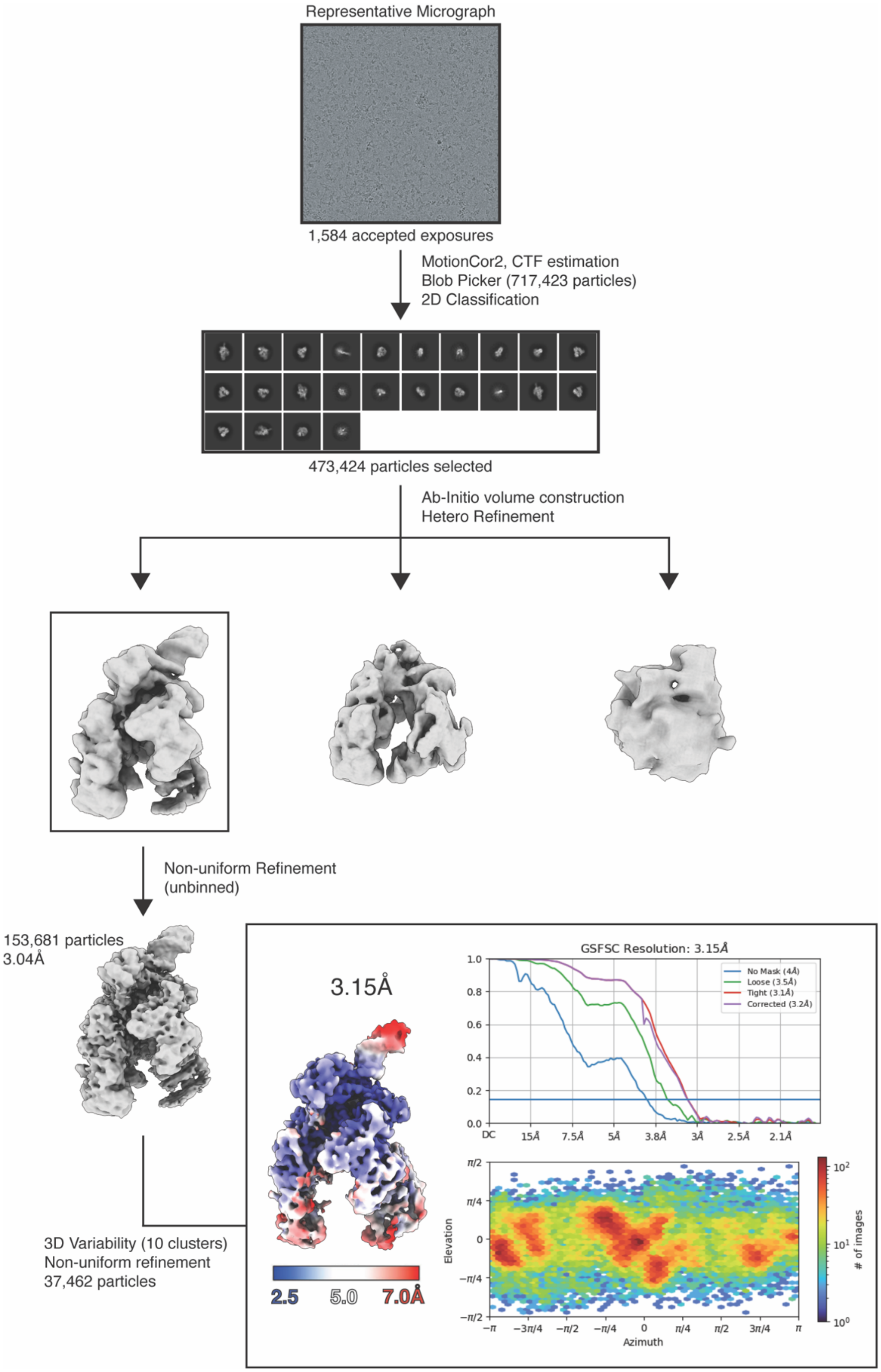


**Supplemental Figure S1**. Cryo-EM workflow for Cas12a-MG29-1. The final sharpened map from cryoSPARC is colored by local resolution. Map used throughout figures was post-processed by EMReady[1]. All processing was performed in CryoSPARC (v4.3) or later versions[2].

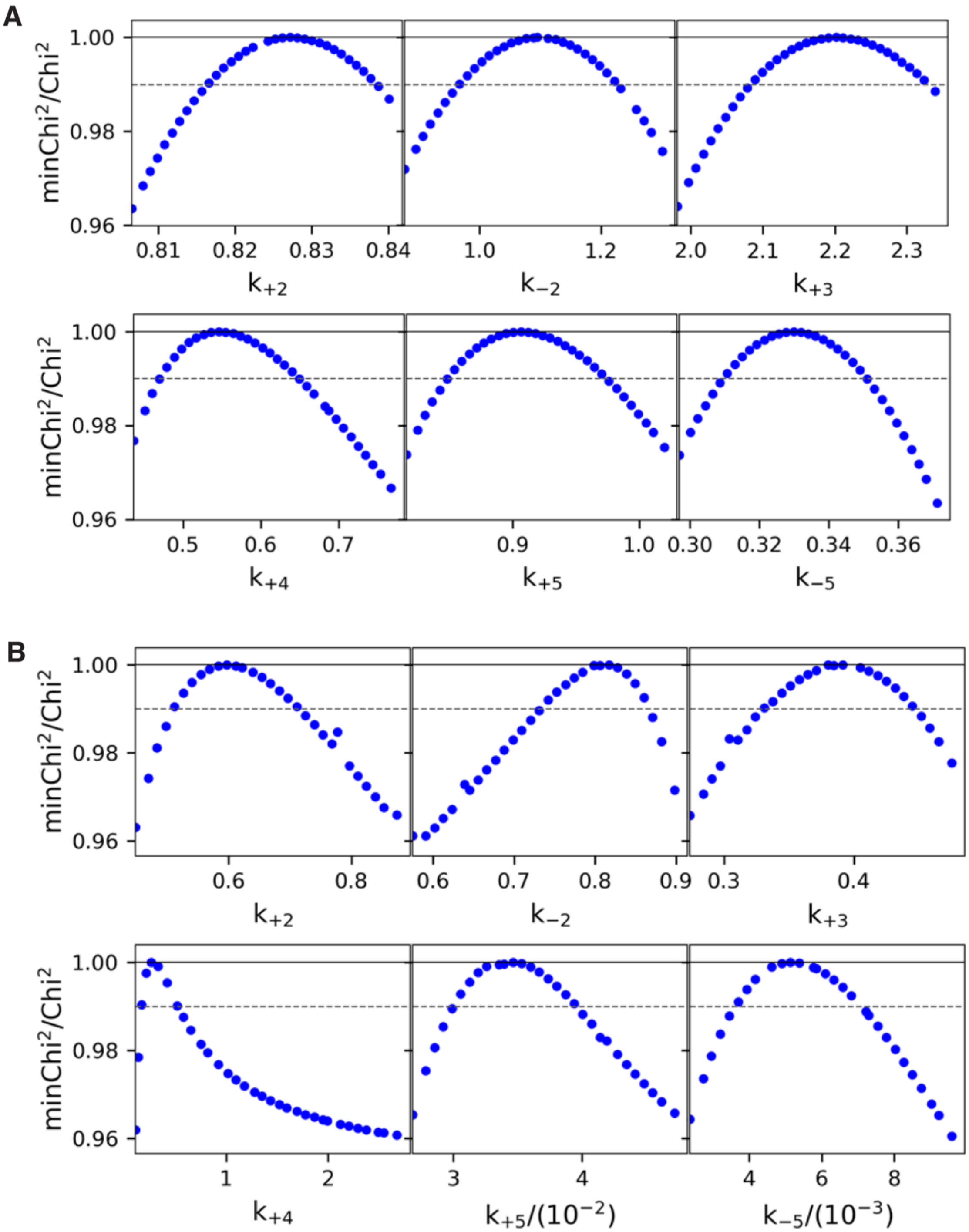


**Supplemental Figure S2**. Confidence contours from global data fitting for Cas12a-MG29-1 in (A) and AsCas12a (B). Each panel displays $\chi^2$ as a function of the corresponding individual rate parameter. The dashed horizontal line denotes the $\chi^2$ value at the 95% confidence level used to derive the parameter limits reported in Supplementary Table 2.

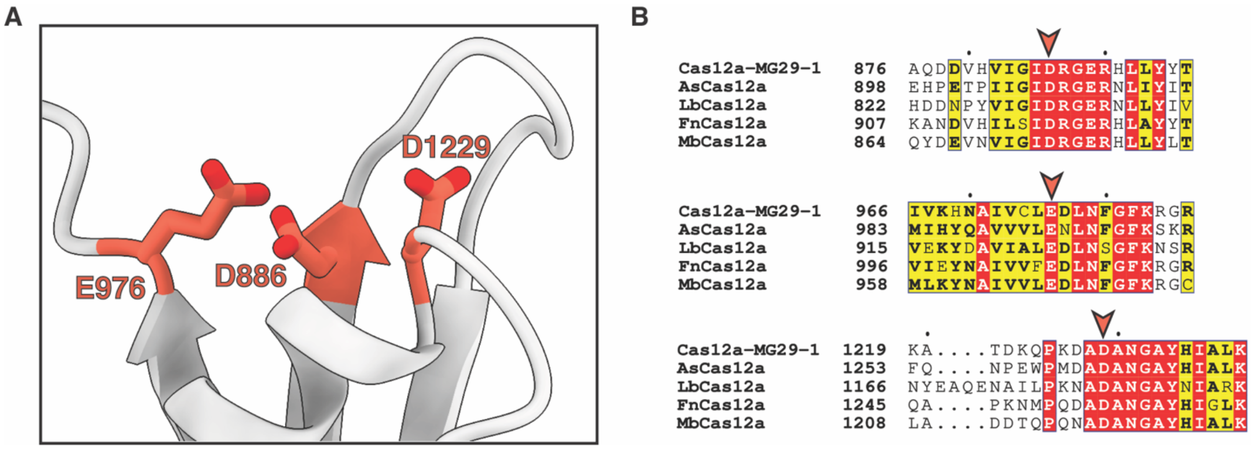


**Supplementary Figure S3.** Active site residues of Cas12a-MG29-1. (A) Putative active site residues D886, E976, and D1229 are colored in orange. (B) Alignment of Cas12a-MG29-1 and Cas12a orthologs near the putative active site residues. The orange pointer indicates each of the conserved active site residues within the alignment. Generated with ESPript.[3]

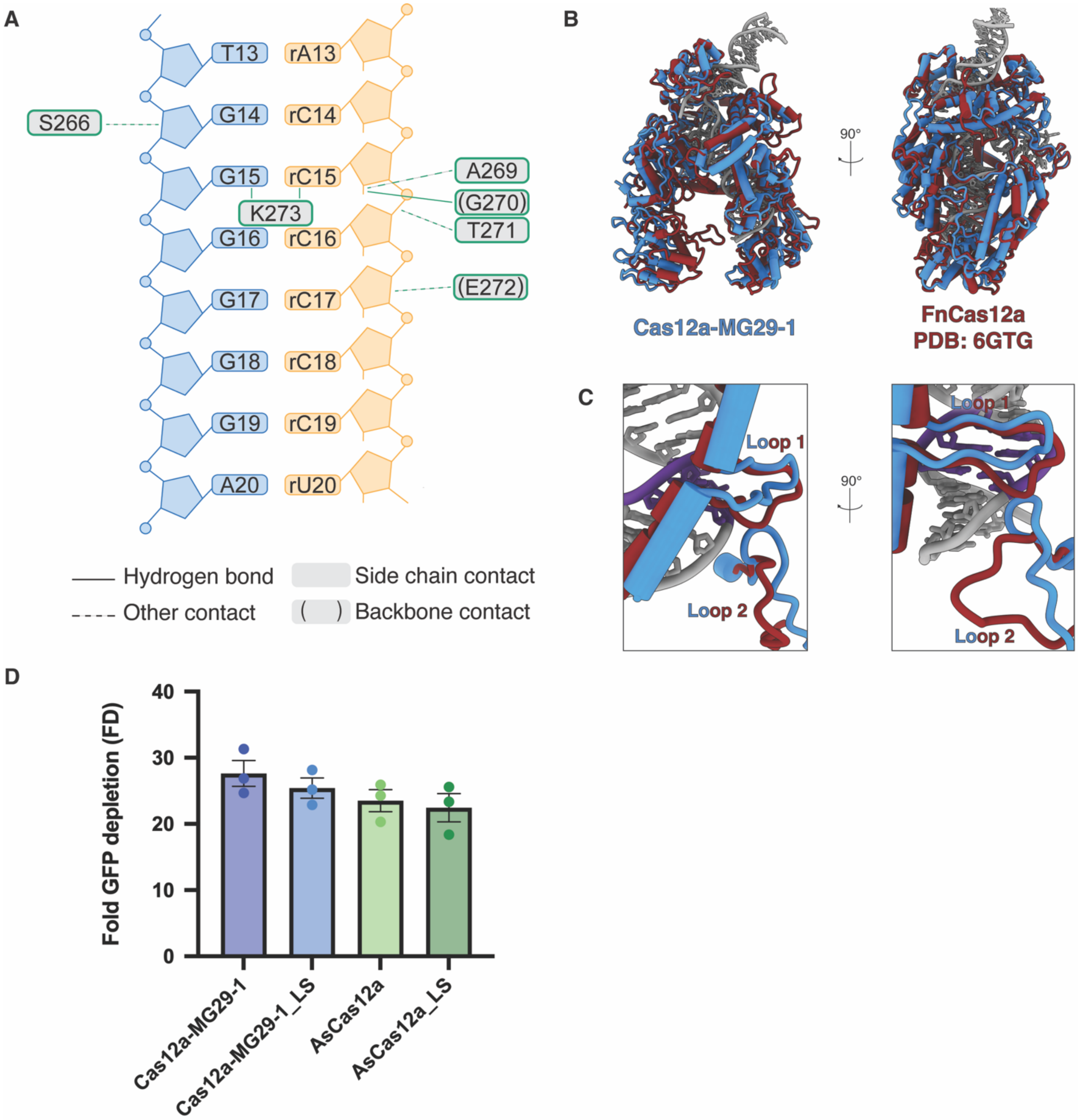


**Supplementary Figure S4.** Structural context of distal R-loop contacts and functional effects of loop swaps. (A) Schematic diagram of Loop 1 nucleic acid-protein contacts within the distal R-loop of AsCas12a.[4] Nucleic acid is colored as follows: crRNA, dark orange; target strand, royal blue. Solid lines indicate hydrogen bonds, while dotted lines denote other contacts. Residues in parentheses refer to side backbone contacts. (B) Structural overlays of the Cas12a-MG29-1 model with FnCas12a (6GTG).[5] Cas12a-MG29-1 is shown in blue, while FnCas12a is shown in brick red. The R-loop is colored gray. Front and side views are shown. (C) Close-up view of Loop 1 and Loop 2. Positions 14-16 of the heteroduplex are highlighted in purple, while the rest of the figure matches the color scheme in (A). Front and side views are shown. (D) Fold GFP depletion of Cas12a-MG29-1, AsCas12a, and loop swap mutants, showing that loop exchange does not substantially alter on-target activity.

### Supplementary Tables

| Oligo | Sequence (5' to 3' ) | Source |
|---|---|---|
| C3_Target_NTS (used for cryo) | CTCACTCCTTTCATTTGGGCAGCTCCCCTACCCCCCTTACCTCTCTAGTC | IDT |
| C3_Target_TS (used for cryo) | GACTAGAGAGGTAAGGGGGGTAGGGGAGCTGCCCAAATGAAAGGAGTGAG | IDT |
| C3_Target_29-1_crRNA (used for cryo) | mG*mU*mU*rGrArGrArArUrCrGrArArArGrArUrUrCrUrCrArArCrCrUrUrUrUrArArU rUrUrCrUrArCrUrGrUrUrGrUrArGrArUrGrGrCrArGrCrUrCrCrCrCrUrArCrCrCrCrCr C*mU | IDT |
| B2MH4_Target_NTS | CTCACTCCTTTCATTTGTCCCGACCCTCCCGTCGCCGTACCTCTCTAGTC | IDT |
| B2MH4_Target_TS | GACTAGAGAGGTACGGCGACGGGAGGGTCGGGACAAATGAAAGGAGTGAG | IDT |
| B2MH4_Target_NTS_FAM | /56-FAM/ CTCACTCCTTTCATTTGTCCCGACCCTCCCGTCGCCGTACCTCTCTAGTC | IDT |
| B2MH4_Target_TS_FAM | /56-FAM/ GACTAGAGAGGTACGGCGACGGGAGGGTCGGGACAAATGAAAGGAGTGAG | IDT |
| B2MH4_Target_NTS_16tCo | CTCACTCCTTTCATTTGTCCCGACCCTCCCGT***tCo***GCCGTACCTCTCTAGTC | Biosynthesis |
| B2MH4_Target_29-1_crRNA | rGrUrUrGrArGrArArUrCrGrArArArGrArUrUrCrUrCrArArCrCrUrUrUrUrArArUrUrUr CrUrArCrUrGrUrUrGrUrArGrArUrUrCrCrCrGrArCrCrCrUrCrCrCrGrUrCrGrCrCrG | IDT |
| B2MH4_Target_As_crRNA | rUrArArUrUrUrCrUrArCrUrCrUrUrGrUrArGrArUrUrCrCrCrGrArCrCrCrUrCrCrCrGr UrCrGrCrCrG | IDT |

**Supplementary Table S1.** Oligonucleotide sequences used in this study. RNA bases preceded by “m” are 2'-O-methyl modified. Asterisks correspond to phosphorothioate linkages.

| Parameter | Cas12a-MG29-1 | AsCas12a |
|---|---|---|
| $k_1$ | 500 μM$^{-1}$s$^{-1}$ * | 500 μM$^{-1}$s$^{-1}$ * |
| $k_{-1}$ | 0.01 s$^{-1}$ * | 0.01 s$^{-1}$ * |
| $k_2$ | 0.83 s$^{-1}$ {0.817,0.838} | 0.6 s$^{-1}$ {0.512, 0.718} |
| $k_{-2}$ | 1.1 s$^{-1}$ {0.967, 1.22} | 0.8 s$^{-1}$ {0.739, 0.86} |
| $k_3$ | 2.2 s$^{-1}$ {2.09, 2.32} | 0.4 s$^{-1}$ {0.338, 0.441} |
| $k_4$ | 0.5 s$^{-1}$ {0.479, 0.64} | 0.3 s$^{-1}$ {0.214, 0.419} |
| $k_5$ | 0.9 s$^{-1}$ {0.854, 0.97} | 0.034 s$^{-1}$ {0.0306, 0.0394} |
| $k_{-5}$ | 0.33 s$^{-1}$ {0.311, 0.349} | 0.005 s$^{-1}$ {0.00369, 0.00682} |
| EDR scaling factor | 3.9 | 2.4 |
| $EDP_1$ | 1.4 | 1.4 |
| $EDP_1P_2$ | 1.4 | 1.4 |

**Supplementary Table S2.** Best fit rate constants and scaling factors from global kinetic fit using the model in Figure 4B. Values in curvy brackets are lower and upper limits based on a 95% confidence interval. An asterisk is used to indicate that DNA binding and dissociation rates were not well constrained by the data, so they were locked to give a $K_d$ of 20 pM.

| | |
|---|---|
| **Data collection** | |
| Microscope | FEI Glacios |
| Voltage (kV) | 200 |
| Detector | Falcon 4 |
| Pixel size (Å/pix) | 0.94 |
| Exposure ($e^-/Å^2$) | 49 |
| Defocus range ($\mu$m) | -1.5 to -2.5 |
| Tilt angle (°) | 0 |
| Micrographs collected | 1,689 |
| Micrographs used | 1,584 |
| Total particles extracted | 717,423 |
| Automation software | SerialEM |
| Sample | Cas12a-MG29-1 + crRNA + DNA target |
| **3D reconstruction statistics** | |
| Particles | 37,362 |
| Symmetry | C1 |
| Map resolution (Å) | 3.15 |
| FSC Threshold | 0.143 |
| Map sharpening B-factor | 76.2 |
| **Model refinement and validation statistics** | |
| Refinement software | PHENIX |
| Initial models | AlphaFold2 (Cas12a-MG29-1), 8SFO (nucleic acid) |
| Composition | |
| Non-hydrogen atoms | 12129 |
| Amino acid residues | 1243 |
| Nucleotides | 91 |
| RMSD bonds (Å) | 0.004 |
| RMSD angles (°) | 0.889 |
| Average B-factors | |
| Amino acids | 81.02 |
| Nucleotides | 62.71 |
| Ramachandran | |
| Favored (%) | 97.65 |
| Allowed (%) | 2.35 |
| Outliers (%) | 0 |
| MolProbity score | 1.22 |
| Clash score | 3.58 |
| Rotamer outliers (%) | 0.91 |
| C-beta outliers (%) | 0 |
| CaBLAM outliers (%) | 0.9 |
| CC (mask) | 0.87 |

**Supplementary Table S3.** Cryo-EM data collection, refinement, and validation statistics for the ternary Cas12a-MG29-1 model.

**SUPPLEMENTAL FIGURES REFERENCES**


1. He, J., Li, T. & Huang, S.-Y. Improvement of cryo-EM maps by simultaneous local and non-local deep learning. *Nat Commun* **14**, 3217 (2023).
2. Punjani, A. Real-time cryo-EM structure determination. *Microanal* **27**, 1156–1157 (2021).
3. Gouet, P. & Courcelle, E. ENDscript: a workflow to display sequence and structure information. *Bioinformatics* **18**, 767–768 (2002).
4. Strohkendl, I. *et al.* Cas12a domain flexibility guides R-loop formation and forces RuvC resetting. *Molecular Cell* **84**, 2717-2731.e6 (2024).
5. Stella, S. *et al.* Conformational Activation Promotes CRISPR-Cas12a Catalysis and Resetting of the Endonuclease Activity. *Cell* **175**, 1856-1871.e21 (2018).
